# GABA administration limits viral replication and pneumonitis in a mouse model of COVID-19

**DOI:** 10.1101/2021.02.09.430446

**Authors:** Jide Tian, Blake Middleton, Daniel L. Kaufman

## Abstract

Despite the availability of vaccines for COVID-19, serious illness and death induced by coronavirus infection will remain a global health burden because of vaccination hesitancy, possible virus mutations, and the appearance of novel coronaviruses. Accordingly, there is a need for new approaches to limit severe illness stemming from coronavirus infections. Cells of the immune system and lung epithelia express receptors for GABA (GABA-Rs), a widely used neurotransmitter within the CNS. GABA-R agonists have anti-inflammatory effects and can limit acute lung injury. We previously showed that GABA treatment effectively reduced disease severity and death rates in mice following infection with a coronavirus (MHV-1) which provides a potentially lethal model of COVID-19. Here, we report that GABA treatment also reduced viral load in the lungs, suggesting that GABA-Rs may provide a new druggable target to limit pulmonary coronavirus replication. Histopathological analysis revealed that GABA treatment reduced lung inflammatory infiltrates and damages. Since GABA is safe for human consumption, inexpensive, and available worldwide, it is a promising candidate to help treat COVID-19.

## Introduction

While GABA-Rs are well known for their role in neurotransmission in the CNS, these receptors are also found on some cells in the periphery, most notably for our studies, on some cells of the immune system and lung epithelia. The biological roles of GABA-Rs on immune cells are not yet well understood, but there is a growing body of evidence that the activation of these receptors has immunoregulatory actions. Rodent and human macrophages and dendric cells express GABA-Rs and GABA-R agonists inhibit their inflammatory activities (1–6). T cells also express GABA-Rs (4–8). The administration of GABA-R agonists inhibits autoreactive Th1 and Th17 cells while promoting CD4^+^ and CD8^+^ Treg responses (5, 6, 9) and ameliorates autoimmune disease in mouse models of type 1 diabetes (T1D), multiple sclerosis, and rheumatoid arthritis, and also limits inflammation in murine type 2 diabetes (1, 5, 6, 10, 11). Human immune cells also express GABA-Rs and GABA inhibits secretion of IL-6, TNF, IL-17A, CXCL10/IP-10, CCL4, CCL20, and MCP-3 from anti-CD3 stimulated PBMC from T1D patients (7). The ability of GABA-R agonists to inhibit the production of number of inflammatory factors is of potential interest for helping to treat COVID-19 since high levels of some of these inflammatory factors in patient sera is associated with the development of severe COVID-19 (12–15).

Lung epithelial cells also express GABA-Rs, specifically the GABA_A_-R subtype. GABA and GABA_A_-R positive allosteric modulators (PAMs) have been shown to reduce inflammation and improve alveolar fluid clearance and lung functional recovery in different rodent models of acute lung injury (16–23), as well as limit pulmonary inflammatory responses and improve clinical outcomes in ventilated human patients (24–26). GABA_A_-R PAMs reduce macrophage infiltrates and inflammatory cytokine levels in bronchoalveolar lavage fluid (BALF) and limit inflammatory responses by rodent and human macrophages (3, 22, 27–31). GABA application reduces the secretion of inflammatory factors from LPS-stimulated human bronchial epithelial cells *in vitro* (17). Finally, GABA can inhibit platelet aggregation (32), which is important because pulmonary thrombosis often occurs in critically ill COVID-19 patients (33, 34).

Mouse hepatitis virus (MHV)-1 is a pneumotropic beta-coronavirus that is widely used as a safe model of SARS-CoV and SARS-CoV-2 infection (35–38). Intranasal inoculation with ≥5 × 10^3^ plaque-forming units (PFU) of MHV-1 in A/J mice induces acute pneumonitis and acute respiratory distress syndrome with a high lethality rate. The MHV-1-infected mice develop clinical symptoms and pathological features similar to those in severely ill COVID-19 patients, including high levels of pulmonary edema, pneumonitis, dense macrophage infiltrates, hyaline membranes, fibrin deposits, accompanied by loss of body weight and respiratory distress (35–38). We previously showed that GABA treatment just after MHV-1 inoculation, or after the appearance of disease symptoms, very effectively protected the mice from severe illness and death (39). Here, we report that GABA treatment also reduced viral loads and pathological findings in their lungs. We discuss possible mechanisms underlying these observations.

## Materials and methods

### Mice

Female A/J mice (8 weeks in age) were purchased from the Jackson Laboratory and maintained in microisolator cages and fed with a standard diet and water *ad libitum*. This study was carried out in accordance with the recommendations of the Guide for the Care and Use of Laboratory Animals of the National Institutes of Health. The protocols for all experiments using vertebrate animals were approved by the Animal Research Committee at UCLA (approval # ARC-2020-122) and were carried out in compliance with the ARRIVE guidelines.

### Reagents

GABA (stock #A2129) was purchased from Millipore-Sigma (St. Louis, MO, USA).

### Virus

MHV-1, DBT cells, and HeLa-CECAM1 were generously provided by Dr. Stanley Perlman (University of Iowa). MHV-I virus was prepared and titered as previously described (35–39).

### Viral infection and GABA treatment

At 9 weeks in age, individual A/J mice were anesthetized and inoculated intranasally with 5 × 10^3^ PFU MHV-1 in 50 μl cold Dulbecco’s modified Eagle’s medium (DMEM). The mice were immediately randomized and provided with plain water (controls) or water that contained GABA (20 mg/mL) as per (9, 39) for the entirety of the observation period. The mice treated with GABA on day 0 are referred to as the GABA_0_ group. Some virus-inoculated mice were provided with plain water for 2 days and treated with GABA beginning on day 3 post-inoculation and are referred to as the GABA_3_ group. The mice were euthanized at 3 or 6 days post-infection and their right and left lungs were dissected for measurements of viral loads and histology, respectively.

### Viral titers

Frozen lung samples were dounce-homogenized into 1 mL of ice-cold DMEM with 10% fetal calf serum and homogenized with 1 mm glass beads using a Qiagen TissueLyser-LT at 50 Hz for 6×1 min. The viral titers in the supernatants were determined by endpoint dilution (40) in HeLa-CEACAM1 cells (85% confluent, 5 × 10^4^ cells/well) using the Spearman-Kärber formula (41) to calculate 50% tissue culture infectious dose (TCID_50_).

### Hematoxylin and eosin staining of lung sections

Their left lungs were fixed in 10% neutral buffered formalin and embedded in paraffin. Lung tissue sections (5 μm) were routine-stained with hematoxylin and eosin. Five images from each mouse were captured under a light microscope at 200 x magnification. The degrees of pathological changes were scored, based on the number of hyaline membranes, % of pulmonary areas with obvious inflammatory infiltrates in lung parenchyma, and the % of area with inflammatory consolidation within the total area of the section. The total numbers of hyaline-like membranes with, or without, cell debris or hyaline-like deposition in alveoli of the lung tissue section were scored as 0: none detectable; 1: 1-5; 2: 5-10; 3: >10. The areas of lung inflammation and hemorrhage in one lung section were estimated and the severity of inflammation and hemorrhage in the section was scored as 1: mild, 2: moderate; 3, marked, 4: severe. Accordingly, an inflammatory score in each mouse was obtained by % of lung areas x severity score. The areas of lung consolidation were estimated in the lung section and scored as 1: <10%; 2:11-25%: 3:26-50%; 4:>50%. Finally, the pneumonitis score of individual mice = (% of inflammation areas x severity score) + lung consolidation score + hyaline membrane score with a maximum score of 11.

## Results

A/J mice were inoculated intranasally with MHV-1 (5 × 10^3^ PFU) and then randomized to receive plain water (controls) or water containing GABA (20 mg/mL, as in (39)) for the remainder of the study. Mice from these groups were euthanized 3 or 6 days post-infection and the virial load in their lungs was determined. Concurrently, another group of MHV-1 inoculated mice was given water containing GABA beginning at 3 days post-infection and the viral load in their lungs was determined 6 days post-infection.

At three days post-infection the mean viral load in the lungs of mice given plain water was about 7-fold higher than that in mice given GABA immediately after infection (p<0.5, Fig. 1). Thereafter, the viral load in the lungs declined as expected, and by day 6 post-infection the viral load in the lungs of control mice were about twice that in the lungs of mice given GABA either immediately or 3 days post-infection, although these differences were not statistically significant (Fig. 1). Thus, early GABA treatment reduced viral loads in the lungs of mice.

**Figure 1.**
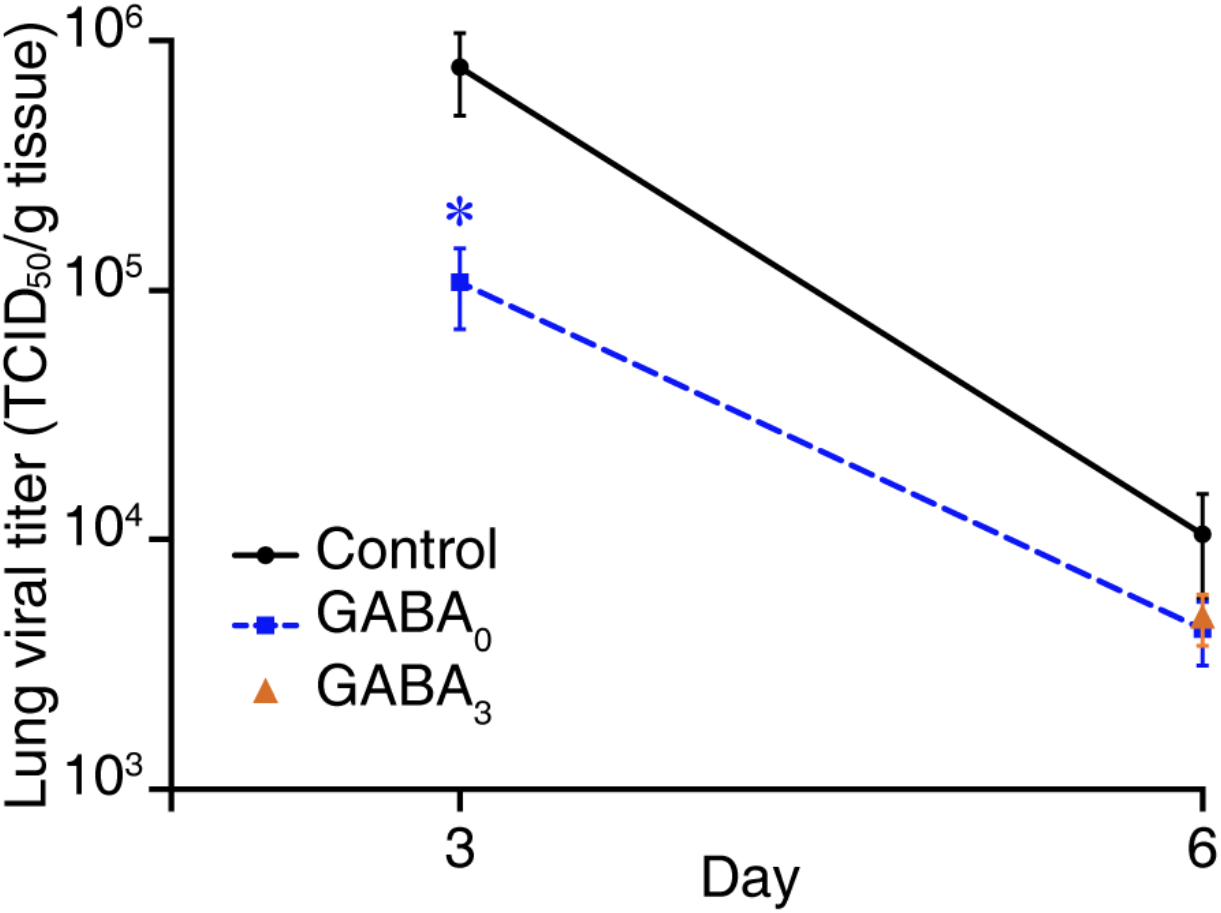
GABA treatment reduces viral replication in MHV-1 infected mice. Mice were inoculated with MHV-1 and immediately placed on plain water (control) or water containing GABA and three or six days later their lungs were harvested for determination of viral load. Concurrently, another group of MHV-1 inoculated mice was given water containing GABA beginning three days post-infection and the viral load in their lungs was determined 6 days post-infection. The data shown are the mean TCID_50_/g of lung tissue ± SEM at the indicated days. GABA_0_ mice (blue square symbol) received GABA immediately after inoculation and GABA_3_ mice (orange triangle symbol) received GABA beginning three days post-infection. N=5 mice per group at each time point. *p<0.05 by Student’s t-test.

Histological evaluation of lung sections from mice given GABA immediately following MHV-1 inoculation revealed reduced inflammatory infiltrates, hyaline-like membrane formation and fibrin deposits in the alveoli at 3 days post-infection, relative to that in control MHV-1 inoculated mice (Fig. 2A). Thus, GABA treatment limited the MHV-1 induced lung damage in A/J mice.

**Figure 2.**
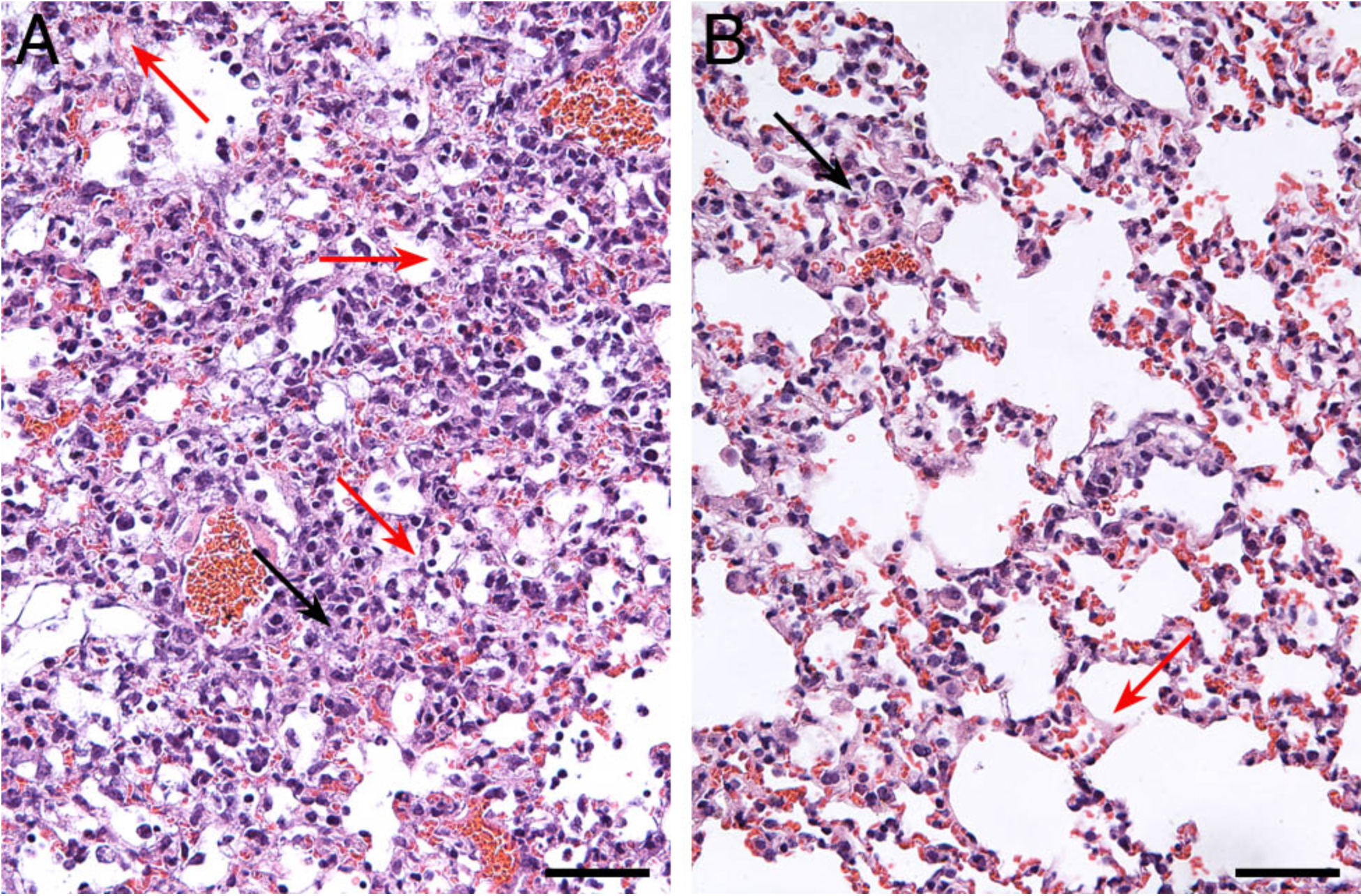
Histopathological features in the lungs of untreated and GABA-treated mice six days post-MHV-1 infection. Images are representative images of H&E stained lung sections from A) untreated mice and B) GABA-treated (beginning immediately following inoculation) mice six days post-infection. Red arrows point to hyaline-like membranes and black arrows indicate local consolidation. Scale bar is 50 um.

## Discussion

Our previous study showed that GABA treatment beginning just after MHV-1 inoculation or after the appearance of symptoms rapidly curtailed disease progression (39). GABA-treated mice also had smaller lung coefficient indexes (indicative of less inflammation and edema). Thus, GABA-R activation can limit a very acute and highly lethal viral infection-induced pulmonary inflammation, a property that heretofore was unknown. Here, we show that early GABA treatment reduced MHV-1 replication in the lungs of mice when measured near the time of peak viral load in this model (35–37), which may have contributed to the better outcomes observed in GABA-treated animals. Hence, GABA-R agonists may provide a new approach to limit viral replication in the lungs.

We can envision a number of ways that GABA may have limited viral replication, including: 1) The lung alveolar cells of rodents and humans express GABA_A_-Rs (30, 42). While the activation of GABA_A_-R’s Cl^−^ channels on neurons leads to Cl- influx and hyperpolarization, the activation of GABA_A_-Rs on ATII cells induces Cl^−^ efflux and greater membrane depolarization (30, 42). Because coronaviruses promote Ca^2+^ influx to enhance their replication (43, 44), the activation of ATII GABA_A_-Rs and the ensuing Cl^−^ efflux and membrane depolarization may limit the influx of extracellular Ca^2+^, making the cellular environment less conducive to viral replication. 2) Activation of GABA_A_-Rs on lung alveolar and large airway epithelial cells may have A) altered the secretion of immune signaling molecules from infected cells, B) altered alveolar surfactant production/absorption, and/or C) altered inflammatory responses and autophagy (45) and, D) reduced the expression of the MHV-1 receptor CAECAM1 in ways that limited virus production. Further detailed studies are needed to evaluate whether these factors, and/or others, contributed to the observed reduction in viral load.

Histopathological analysis of lungs from mice that received GABA just after MHV-1 inoculation revealed that three days post-infection these mice had significantly reduced lung inflammation and markers of lung damage relative to that in control mice. There are a number of different biological processes through which GABA treatment may have ameliorated the severity of pneumonitis: 1) GABA can inhibit macrophage and dendritic cell inflammatory activities (1–3, 29, 46, 47). Likewise, GABA_A_-R PAMs reduce the numbers of macrophages in broncholavage fluid, lung secretion of inflammatory cytokines, and inflammatory responses by rodent and human macrophages (22, 27–31). GABA_A_-R agonists also inhibit activated Th17 and Th1 responses and promote CD4^+^ and CD8^+^ Tregs, however, since adaptive immune responses take time to arise, these abilities are unlikely to have contributed to GABA’s ability to attenuate disease soon after MHV-1 infection. These effects on adaptive immune responses may be relevant for treating COVID-19 which has a longer disease course and in which high levels of circulating Th1, Th17, and Th2-secreted proteins are associated with severe illness (12, 13). 2) By reducing viral loads in the lungs, GABA treatment may have limited dysregulated inflammatory responses to the infection. 3) GABA and GABA_A_-R PAMs reduce inflammation and improve alveolar fluid clearance and lung functional recovery in animal models of acute lung injury (16–18, 20–23) and in ventilated patients (24–26) and could have exerted similar actions in the MHV-1 infected mice. 4) GABA and GABA_A_-R agonist treatments increase macrophage autophagy (45). In murine models of pneumatic bacterial infections, GABA_A_-R agonists reduced bacterial load and TNFα and IL-6 levels in the lungs and protected the mice against illness (45). 5) GABA inhibits platelet aggregation (32)—this may be an important property because pulmonary thrombosis is increased in critically ill COVID-19 patients (33, 34). Thus, treatment with GABA may have led to better outcomes in MHV-1 infected mice through multiple and diverse pathways.

Much remains to be learned about the mechanisms by which GABA-R activation protected MHV-1 infected mice from severe pneumonitis and whether these observations extend to SARS-CoV-2 infections. Given that GABA can affect many biological processes and that viral infection is a very dynamic process it is clear that GABA-R agonist dosing needs to be carefully studied and optimized for different stages of coronavirus infection. Our observations provide a spring board for future investigations into whether the GABA system can be modulated to limit severe illness due to SARS-CoV-2, future novel coronavirus outbreaks, and other respiratory disorders.

## Acknowledgments

We would like to thank Dr. Stanley Perlman for generously providing MHV-1, DBT cells, and HeLa-CECAM1 cells. We would also like Drs. Min Song for assistance. This work was supported by a grant to DLK from the UCLA DGSOM-Broad Stem Cell Research Center COVID-19 Research Award (ORC #20-34) and DLK’s unrestricted funds.

## Disclosures

DLK and JT are inventors of GABA-related patents. DLK serves on the Scientific Advisory Board of Diamyd Medical. BM has no financial conflicts of interest.

## Author Contributions

Conceived and designed the experiments: JT, DLK; Performed the experiments: JT, BM; Analyzed the data: JT, DLK. Wrote the paper: JT, DLK. Drs. Daniel Kaufman and Jide Tian are guarantors of this work and, as such, had full access to all the data in the study and take responsibility for the integrity of the data and the accuracy of the data analysis. All authors approved the final manuscript as submitted.

## Notes

### Summary of Updates

The revision clarifies the legend for Fig. 1.

